# Recombination hotspots in *Neurospora crassa* controlled by idiomorphic sequences and meiotic silencing

**DOI:** 10.1101/2023.11.13.566876

**Authors:** P. Jane Yeadon, Frederick J. Bowring, David E. A. Catcheside

**Affiliations:** College of Science & Engineering, Flinders University, Adelaide, South Australia, AUSTRALIA

## Abstract

Genes regulating recombination in specific chromosomal intervals of *Neurospora crassa* were described in the 1960s but the mechanism is still unknown. For each of the *rec-1*, *rec-2* and *rec-3* genes, a single copy of the putative dominant allele, for example *rec-2SL* found in St Lawrence OR74 A wild type, reduces recombination in chromosomal regions specific to that gene. However, when we sequenced the recessive allele, *rec-2LG* (derived from the Lindegren 1A wild type) we found that a 10 kb region in *rec-2SL* strains was replaced by a 2.7 kb unrelated sequence, making the “alleles” idiomorphs. When we introduced *sad-1*, a mutant lacking the RNA-dependent RNA polymerase that silences unpaired coding regions during meiosis into crosses heterozygous *rec-2SL/rec-2LG*, it increased recombination, indicating that meiotic silencing of a gene promoting recombination is responsible for dominant suppression of recombination. Consistent with this, mutation of *rec-2LG* by RIP (repeat induced mutation) generated an allele with multiple stop codons in the predicted *rec-2* gene, which does not promote recombination and is recessive to *rec-2LG*. *sad-1* also relieves suppression of recombination in relevant target regions, in crosses heterozygous for *rec-1* alleles and in crosses heterozygous for *rec-3* alleles. We conclude that for all three known *rec* genes, one allele appears dominant only because meiotic silencing prevents the product of the active, “recessive”, allele from stimulating recombination during meiosis. In addition, the proposed amino acid sequence of REC-2 indicates regulation of recombination in Neurospora differs from any currently known mechanism.

## Introduction

In eukaryotes, meiotic recombination frequency varies across each genome and between individuals in a species (CATCHESIDE 1975; LINDAHL 1991; ELLEGREN *et al*. 1994). Local recombination is influenced by the position and sequence of recombination hotspots such as *cog^+^* in *Neurospora crassa* (ANGEL *et al*. 1970), the *ade6 M26* mutation in *S. pombe* (GUTZ 1971) and the PRDM9 recognition sequence (BERG *et al*. 2010), a degenerate 13 base motif (CCNCCNTNNCCNC) found in mammalian recombination hotspots (MYERS *et al*. 2008), and probably in most vertebrates (BAKER *et al*. 2017) except birds (SINGHAL *et al*. 2015) and canids (CAMPBELL *et al*. 2016). In *Saccharomyces cerevisiae* (PAN *et al*. 2011) and in other organisms lacking PRDM9, there is a mechanism that targets recombination initiation to regions under substantial selection (BRICK *et al*. 2012; LAM AND KEENEY 2015; SINGHAL *et al*. 2015; CAMPBELL *et al*. 2016), such as promoters, making such hotspots more evolutionarily stable than those specified by PRDM9 (BAKER *et al*. 2017). This seems to be a default mechanism (BAKER *et al*. 2017), as most recombination events in *Prdm9* knockout mice are initiated at promoters, which is rarely the case in wild-type mice (BRICK *et al*. 2012).

The PRDM9 protein (BERG *et al*. 2010) is a SET-domain histone H3 lysine 4 trimethyltransferase with a zinc finger array that binds to the hotspot recognition sequence. This zinc finger array varies between *PRDM9* alleles and, as the motif is also highly polymorphic (JEFFREYS AND NEUMANN 2002; JEFFREYS AND NEUMANN 2005; DURBIN *et al*. 2010), the interaction between *PRDM9* alleles and motifs provides the mechanism for region-specific regulation of recombination (BAUDAT *et al*. 2010; MYERS *et al*. 2010; PARVANOV *et al*. 2010).

Like *PRDM9*, the *N. crassa rec* genes, *rec-1*, *rec-2* and *rec-3*, influence recombination at specific loci (Fig 1), suggesting they encode proteins (CATCHESIDE 1977). *rec-2* is in the 450 kb region between the *sp* and *am* loci on Linkage Group V (SMITH 1966; SMITH 1968; CATCHESIDE AND CORCORAN 1973), and modulates allelic recombination in *his-3*, a histidine biosynthesis gene on the right arm of LGI (PERKINS *et al*. 1982). *rec-2* also affects crossing over in at least three intergenic intervals, including between *his-3* and *ad-3* (Fig 1) (CATCHESIDE 1975; CATCHESIDE 1977). *rec-2* is thought to interact with the recombination hotspot *cog* (ANGEL *et al*. 1970), ~3 kb distal of the 3′ end of *his-3* (BOWRING AND CATCHESIDE 1991; YEADON AND CATCHESIDE 1995), between *his-3* and *ad-3* (CATCHESIDE AND ANGEL 1974). *cog* is naturally polymorphic, with two codominant alleles (ANGEL *et al*. 1970; YEADON *et al*. 2004) of which *cog^+^* stimulates ~25 times more recombination than *cog* (YEADON *et al*. 2004). When *rec-2SL* (previously known as *rec-2^+^*) is present in a cross, recombination is independent of the *cog* genotype (CATCHESIDE AND ANGEL 1974) and is reduced by up to 100-fold (YEADON *et al*. 2004; YEADON *et al*. 2016).

**Fig 1.**
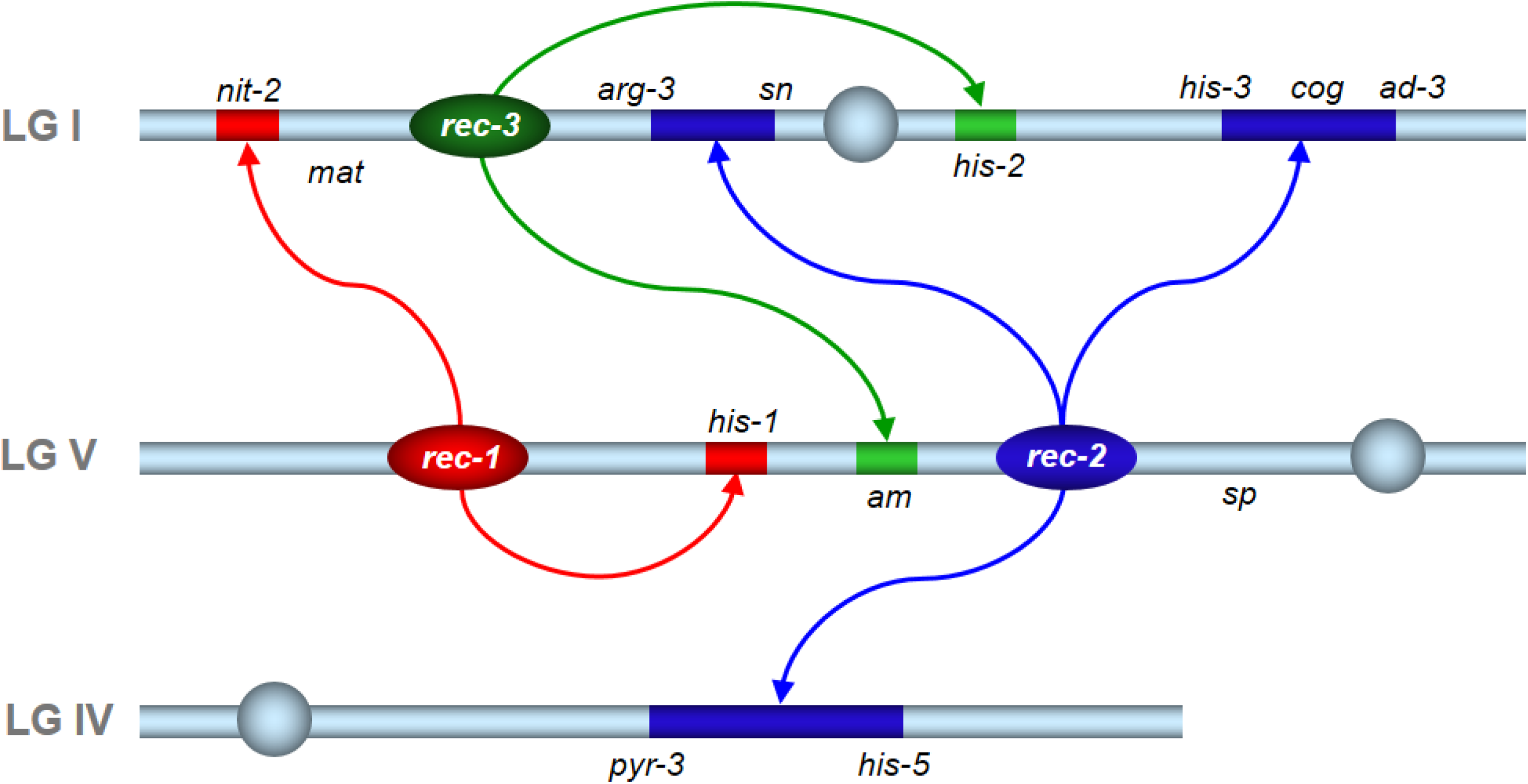
Neurospora *rec* genes and areas of influence. The orientation of LGV is inverted in the diagram to ease illustration. The colour-coded arrows indicate chromosomal regions influenced by each similarly coloured *rec* gene. Based on (CATCHESIDE 1977).

Both *rec-1SL* and *rec-3SL* (previously known as *rec-1^+^*and *rec-3^+^* respectively) appear to act in a similar way to *rec-2SL*, with the dominant allele reducing recombination in the target region of each gene (JESSOP AND CATCHESIDE 1965; CATCHESIDE 1966). Of recombination gene/recognition-site systems, including not only PRDM9 but also recBCD/*Chi* in *E. coli* (PONTICELLI *et al*. 1985; AMUNDSEN *et al*. 2016) and the *HO* endonuclease for mating type switching in *S. cerevisiae* (NICKOLOFF *et al*. 1986), Neurospora *rec* genes appeared unique for their dominant reduction of recombination. The Neurospora St Lawrence wild type strain used for the initial genome sequence (GALAGAN *et al*. 2003), 74-OR23-1VA (FGSC#2489), carries all three of the dominant *rec^+^* alleles, which at first seemed a happy coincidence. On the contrary, the discovery of meiotic silencing (SHIU *et al*. 2001) showed that the choice of 74-OR23-1VA was very unfortunate for those interested in *rec* genes, as it has now been shown not to contain the active versions of any of the *rec* genes (this manuscript).

Several mechanisms to defend against proliferation of transposable elements exist in Neurospora (BILLMYRE *et al*. 2013), including repeat-induced point mutation (RIP; (SELKER 1990a)), which occurs premeiotically, after fertilization but before nuclear fusion. If a sequence exists in more than a single copy in a haploid nucleus, both copies of duplicated sequences experience high frequency GC→ AT mutations (CAMBARERI *et al*. 1989).

Meiotic silencing by unpaired DNA (SHIU *et al*. 2001), another Neurospora defence mechanism, is a system of RNA silencing similar to co-suppression in plants (NAPOLI *et al*. 1990), RNAi in animals (FIRE *et al*. 1998) and quelling in fungi (COGONI *et al*. 1996), all of which require RNA-dependent RNA polymerases (reviewed in (BILLMYRE *et al*. 2013)). Meiosis is the only diploid phase in the predominantly haploid *N. crassa* life cycle. If a sequence is unpaired during meiosis, not only is the unpaired transcript silenced but also any additional homologous transcripts, whether paired or not (SHIU *et al*. 2001), although silencing is often incomplete (SHIU *et al*. 2001). Thus, for a foreign gene inserted into the Neurospora genome, or an endogenous Neurospora gene where one copy is present, there may be insufficient protein product for normal function during meiosis.

Knowledge of meiotic silencing (SHIU *et al*. 2001) allowed us to determine that some *rec* alleles appear dominant not because their products are active but because the sequence lacks homology to the sequence at the same location on the homologous chromosome, the product of which is active and required to stimulate recombination. The lack of a pairing partner silences expression of the active sequence in a heterozygote. In addition, we have shown that Neurospora *rec* genes provide an additional example to PRDM9 of regulation of local meiotic recombination.

## Materials and Methods

### Data and Reagents

Neurospora strains are available on request. Sequence data are available at GenBank; the accession numbers are listed in the manuscript.

### Fungal culturing and media

Neurospora media and culture methods were as described previously (BOWRING AND CATCHESIDE 1996). Recombination assays and crosses were as described previously (YEADON *et al*. 2004; YEADON *et al*. 2010). In Table S1, for *his-3*^K874^ × *his-3*^K1201^ crosses, spore suspensions used for selective counts (plates lacking histidine) were diluted 800-fold for viable counts (fully supplemented plates), while for crosses between *am* or *his-1* alleles, suspensions used for selective counts were diluted 400-fold for viable counts.

### Construction of Neurospora strains

*Sad-1 rec-2SL his-3*^K874^ strains (T12739 and T12741; Table S2) were extracted from a cross between T12322 and T10989, as were T12740 and T12742. *Sad-1 rec-1SL his-1*^K83^ strains (T12397 and T12398) were extracted from a cross between T12322 and T3881. Further *his-1* strains (T12815-T12820) were progeny of T12398 × F6207. *Sad-1 rec-3SL* strain T12394 was extracted from a cross between T1826 and T12322. *Sad-1 rec-3SL am*^47305^ strains (T12429 and T23430) were extracted from a cross between T12394 and T813, as were T12427 and T12428.

*rec-2LG* deletion constructs, containing sequences flanking the *rec-2LG* region surrounding *hph*, were introduced by electroporation (MARGOLIN *et al*. 1997) into T10998, to give T12821, in which *rec-2LG* was replaced with *hph*. Further *rec-2* deletion strains (T12823-T12831) were extracted from a cross between T12821 and T11805.

*rec-2LG* replacement constructs, containing sequence covering the *rec-2LG* region and *hph*, were introduced by electroporation (MARGOLIN *et al*. 1997) into *rec-2SL* strains T12564 and T10997, *his-3*^K480^ and *his-3*^K874^ respectively. Homokaryotic Hyg^R^ transformants were isolated and subjected to Southern analysis. Transformants in which there seemed to be a single insertion event, replacing the *rec-2SL* sequence with that of *rec-2LG* and the *hph* (Hyg^R^) gene, were crossed to *cog^+^ rec-2LG* strains T10998 or T11805, *his-3*^K874^ and *his-3*^K1201^ respectively. His^+^ frequencies in the crosses of T12564 transformants (T12744–T12747; Table S2) to T10998 were all in the expected range for *rec-2SL* crosses (Table S3), but of the crosses between T10997 transformants and T11805, T12749 × T11805 yielded a 20-fold higher His^+^ frequency (Table S3). After extraction of Hyg^R^ *am* progeny from this cross (T12755–T12760; Table S2), a cross between T12757 and T12760 increased His^+^ frequency a further 4-fold (Table S3). In progeny extracted from T12757 × T12760 (T12762-T12771), it was clear that two alleles of *rec-2, rec-2^HF^* and *rec-2^RIP^*, were segregating (Table S4).

### Locating crossovers either side of *rec-2*

Since *rec-2SL* lies between *sp* and *am*, *sp^+^ am^+^* progeny from cross B539 (BOWRING AND CATCHESIDE 1999) heterozygous *rec-2LG/rec-2SL*, with *am^+^*and *sp^+^* markers *in trans*, were selected, thus resulting in strains that had experienced crossing over on one or the other side of *rec-2SL*. Using cosmid (ORBACH 1994) probes and Southern analysis, restriction site polymorphisms (RSPs) flanking each crossover position were identified (BOWRING AND CATCHESIDE 1996; BOWRING AND CATCHESIDE 1999). When sequence covering the *rec-2SL* region became available (AIGN *et al*. 2001), additional RSPs were identified within PCR amplicons, using primer pairs A2F/A2R…BE17F/BE17R (Table S5; Fig 2). A combination of PCR-based RSP markers, A2, B, C and E, and markers determined by presence/absence of amplicons were used to further refine the position of *rec-2SL* by locating crossovers in additional recombinant *sp^+^ am^+^* progeny from cross B539.

**Fig 2.**
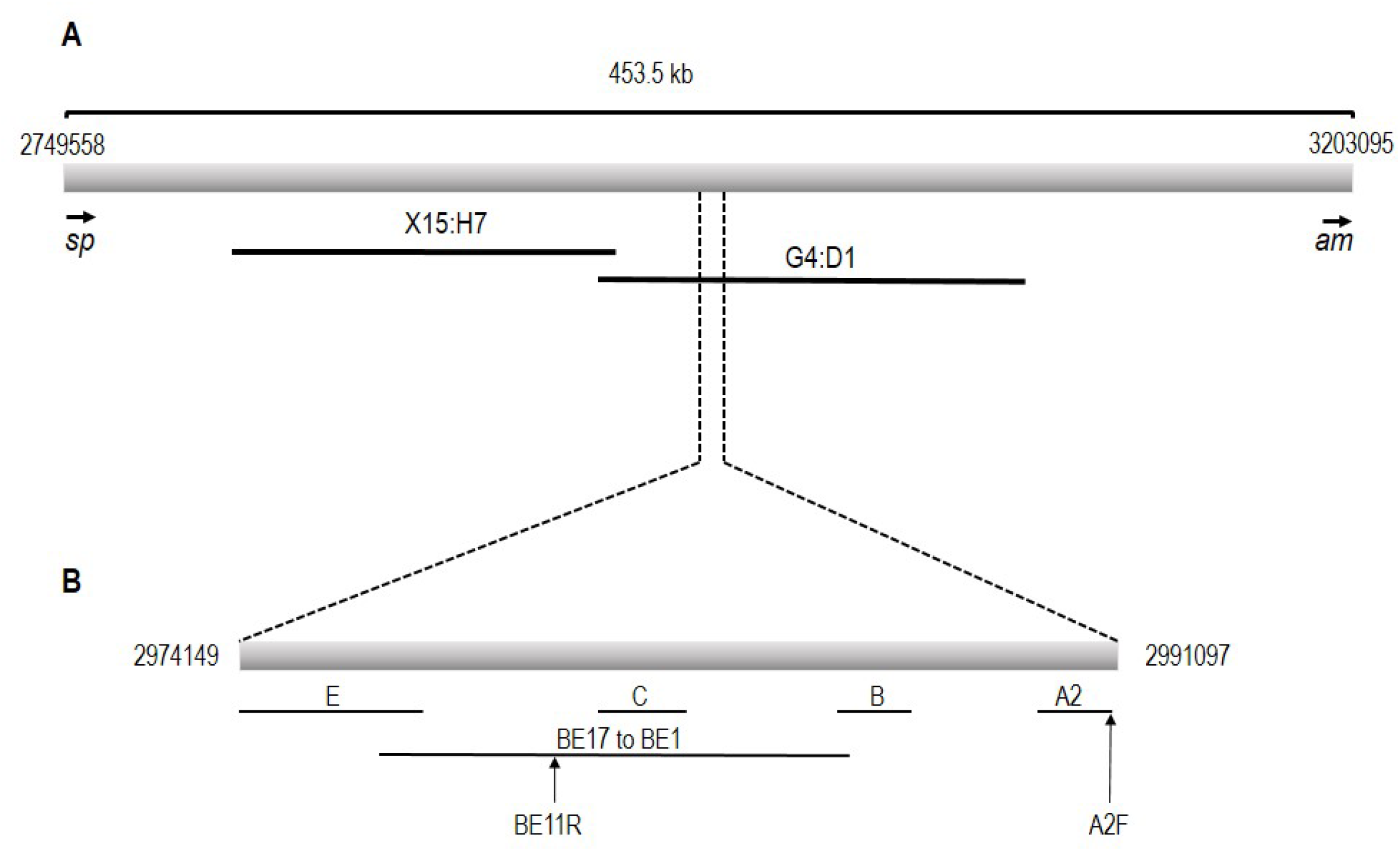
Markers used to locate the *rec-2 SL/rec-2LG* regions within the *am* to *sp* interval. **A.** The LGV centromere is to the left of the figure. The locations of cosmids X15:H7 and G4:D1 (ORBACH 1994) are shown below the 69 kb region they cover. **B.** The region covered by BE1 to BE17 is shown as a single bar. Bars labelled A2, B, C and E indicate the positions of markers used in *rec-2* mapping. A2F and BE11R are primers located at the right end of A2 and the left end of BE11 respectively, yielding amplicon A2F/BE11R. Positions are those in NC12 of the *N. crassa* OR74A genome assembly.

### Generation of *rec-2* replacement DNA constructs

Left component (LF) includes 727 bases of sequences common to both *rec-2LG* and *rec-2SL*, extending into the 2691 bp *rec-2LG*-specific region. *rec-2LG* genomic DNA from strain T10998 was amplified by PCR using primers A2F and rec2loxR (Table S5), and this amplicon was subsequently fused to “HY”, the most 5ʹ 1101 bases of the *hph* gene that encodes hygromycin-B-phosphotransferase (GRITZ AND DAVIES 1983).

In the right component (RF), the 3ʹ terminal 1086 bases of *hph*, “YG”, which has a 657 bp overlap with the 5ʹ component of *hph*, is fused to 1007 bases of sequence common to *rec-2LG* and *rec-2SL* strains PCR-amplified using primers 3rec2loxF and 3rec2R (Table S5).

For deletion constructs, the RF is the same as for replacement constructs, but in the left component, LF is fused directly to HY.

### Integration of constructs

Transformation was by electroporation as described previously (MARGOLIN *et al*. 1997). Selection of Hyg^R^ transformants ensured that both fragments are inserted in the chromosome in such a way that “HY” and “YG” recombined to give a functional *hph* gene, a “split marker” transformation (CATLETT *et al*. 2003). To improve the otherwise poor yield in *Neurospora* of the desired construct, the left and right flank sections include respectively 727 and 1007 bases shared between *rec-2LG* and *rec-2SL*.

### Generation of *rec-2LG* sequence and *rec-2^RIP^* sequences

For the *rec-2LG* sequence, PCR was performed using genomic DNA from a *rec-2LG* strain as template and primers A2_2F and BE11R. Internal primers were generated from each round of sequencing, using initially the amplification primers as sequence primers. These sequencing primers are listed in the last eight positions in Table S5.

For mutated *rec-2* sequence, genomic DNA from *rec-2^HF^* strain T12762 and *rec-2^RIP^* strain T12763 (Table S2) were used as templates. rec-2delLF and HY, which are inside the insertion sequence, were used as primers (Table S5). The subsequent amplicon was sequenced using the sequencing primers used to obtain *rec-2LG* sequence.

## Results

### Location of *rec-2SL*

Restriction site polymorphisms (RSP) within the ~450 kb *sp*–*am* interval on LGV were used to locate crossovers on either side of *rec-2SL* (Fig 2) in 238 *sp^+^ am^+^* progeny from a cross *sp rec-2LG am^+^* × *sp^+^ rec-2SL am* (BOWRING AND CATCHESIDE 1999). As a result, *rec-2SL* was found to lie within a 69 kb region (Fig 2) covered by two pMOcosX cosmids (ORBACH 1994), G4:D1 and X15:H7. RSP markers in the A2 to E region (Fig 2) were used to show that the *rec-2LG*/*rec-2SL* difference falls in the 13107 bp interval between markers B and E (Fig 2). However, *rec-2LG* DNA in the BE1 to BE11 region (Fig 2) could not be amplified by PCR despite success with *rec-2SL* DNA. In addition, using A2F and BE11R as PCR primers (Fig 2) yielded amplicons that measured ~11.1 kb and ~3.5 kb respectively using *rec-2SL* and *rec-2LG* DNA as template, demonstrating that the difference between *rec-2SL* and *rec-2LG* falls within an insertion/deletion region.

### *rec-2SL* and *rec-2LG* are idiomorphs

Since unpaired DNA is silenced during meiosis in *N. crassa* (SHIU *et al*. 2001), if the recessive allele, *rec-2LG*, were to encode an activator of recombination and fall within an insertion/deletion region, it would lack a pairing partner in a *rec-2LG/rec-2SL* heterozygote. Subsequent meiotic silencing could explain both the dominant phenotype of *rec-2SL* and the lack of influence of *rec-2SL*-specific sequence.

*Sad-1* is a dominant mutation in an RNA-dependent RNA polymerase required for meiotic silencing (SHIU AND METZENBERG 2002). Ironically, *Sad-1* is dominant because it is a deletion mutant, and silences itself (SHIU AND METZENBERG 2002). Thus, in crosses carrying the *Sad-1* mutant and one copy of *rec-2SL*, an increase in *his-3* allelic recombination would support the hypothesis that meiotic silencing explains dominance of low frequency recombination in heterozygous crosses. In *Sad-1 rec-2SL* × *rec-2LG* crosses between *his-3*^K874^ and *his-3*^K1201^ alleles, His*^+^* frequency is ~9-fold higher than is typical in *rec-2SL* crosses (Table 1; Table S1), although only ~30% of that typical of crosses homozygous *rec-2LG* (Table S1; (YEADON *et al*. 2004)). The influence of *Sad-1* on recombination at *his-3* is restricted to a *rec-2LG/rec-2SL* heterozygote (Table 1; Table S1). In addition, a deletion of *rec-2LG* (Δ*rec-2*), where the *rec-2LG* region is replaced with the hygromycin-resistance gene sequence, behaves similarly to *rec-2SL* with a low *his-3* allelic recombination frequency when Δ*rec-*2 is heterozygous (Table 2; Table S6). These data confirm the *rec-2LG* recombination activator hypothesis: the activating gene is missing in Δ*rec-2LG* homozygotes while the lack of a pairing partner silences the activating gene in both *rec-2LG/rec-2SL* and *rec-2SL/*Δ*rec-2LG* heterozygotes.

**Table 1.**
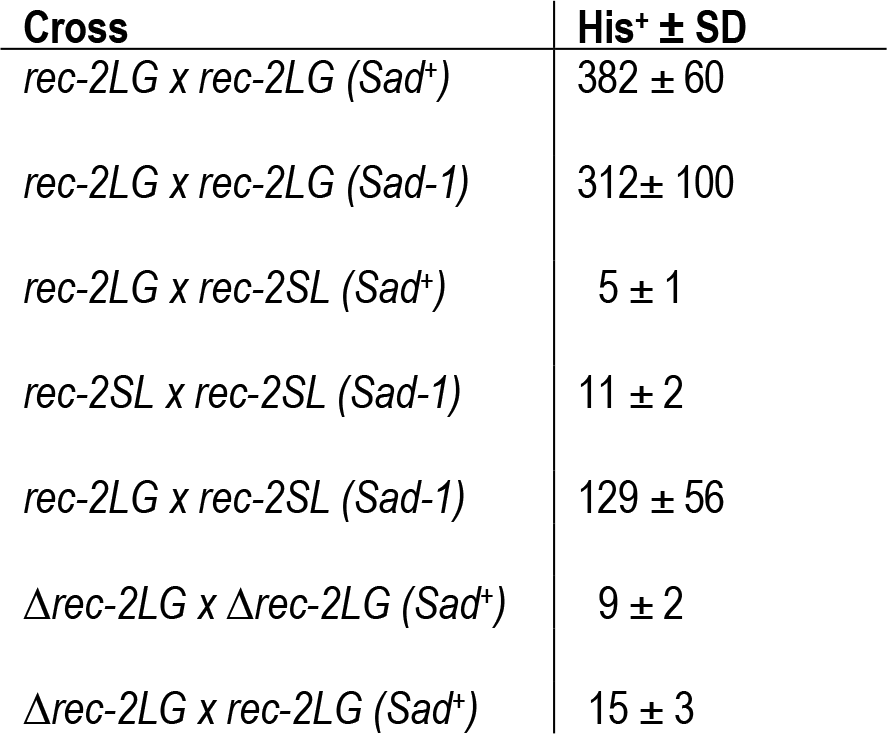
Recombination frequency at *his-3* in crosses between *his-3*^K874^ and *his-3*^K1201^ alleles with and without meiotic silencing. His^+^ ± SD indicates the frequencies of His^+^ progeny per 10^5^ viable spores +/- sample standard deviation. Detailed cross data are provided in Table S1. Neurospora strains used for this work are listed in Table S2.

**Table 2.**
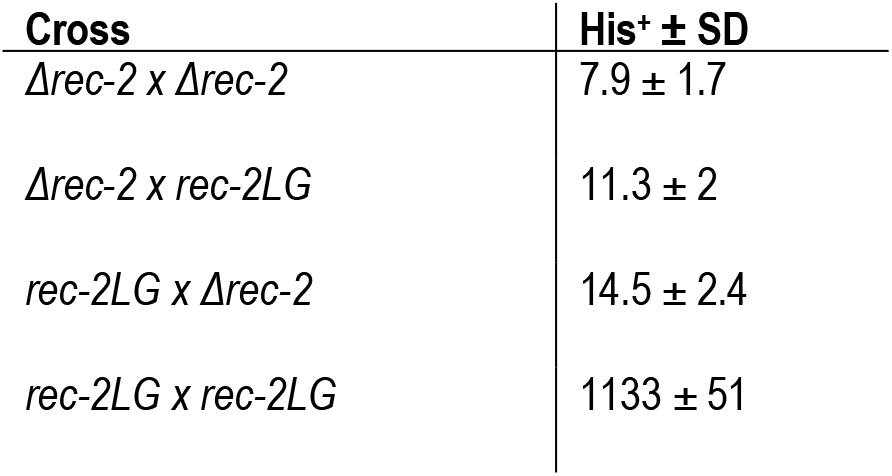
Recombination frequency at *his-3* in crosses between *his-3*^K874^ and *his-3*^K1201^ alleles with and without deletion of *rec-2*. His^+^ ± SD indicates the frequencies of His^+^ progeny per 10^5^ viable spores +/- sample standard deviation. Detailed cross data are provided in Table S6. Neurospora strains used for this work are listed in Table S2.

### Evidence that *rec-1* and *rec-3* “alleles” are also idiomorphs

The St Lawrence *rec-1* and *rec-3* alleles also behave as dominant suppressors of recombination at *his-1* and *am* respectively (JESSOP AND CATCHESIDE 1965; CATCHESIDE 1966). In *his-1*^K83^ × *his-1*^K90^ and *am^47305^* × *am^K314^* crosses in which meiotic silencing is disabled (*Sad-1 rec-1SL* × *rec-1EmA* (from Emerson *A*) and *Sad-1 rec-3SL* × *rec-3Ema* crosses (from Emerson *a*)), we found that recombination frequencies in *recSL* heterozygotes are high, similar to those in *recEmA* or *recEma* homozygotes (Table 3). This indicates that, like *rec-2LG*, *rec-1EmA* and *rec-3Ema* are activators of recombination, and dominance of each *recSL* allele is due to meiotic silencing.

**Table 3.**
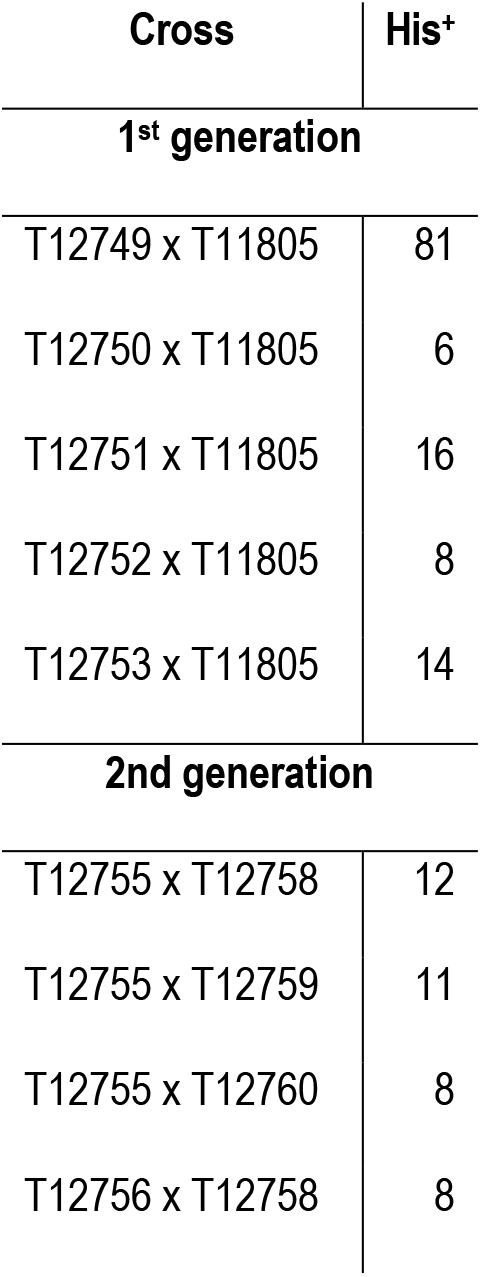
Recombination frequency at *am* and *his-1* in *recSL/rec* crosses with and without meiotic silencing. *am* alleles are *am^K314^* and *am^47305^*; *his-1* alleles are *his-1^K83^* and *his-1^K90^*. RF±SD is prototrophic progeny (with respect to histidine or alanine respectively) per 10^5^ viable spores +/- sample standard deviation. Detailed cross data (#) are in Table S1. Other data (¥, ¶) are historical (JESSOP AND CATCHESIDE 1965; CATCHESIDE 1966). Neurospora strains used for this work are listed in Table S2.

### *rec-2LG* sequence has no similarity to PRDM9

Sequence from the *rec-2LG* A2F-BE11R interval (Fig 2, Table S5) shows that the *rec-2LG* genome lacks 10,234 bases present in the 74-OR23-1VA genome, carrying instead 2,647 bases of *rec-2LG*-specific DNA (Fig 3). The *rec-2LG*-specific sequence begins immediately after nucleotide 2980137 and ends before nucleotide 2990371 in supercontig 12.5 (LGV) in NC12 of the *N. crassa* OR74A genome assembly (GALAGAN *et al*. 2003). Using *N. crassa* parameters, Augustus (STANKE *et al*. 2004) predicts a single open reading frame (ORF) with two introns (cds#1 in Fig 3, Fig S7), resulting in a 762 amino acid protein.

**Fig 3.**
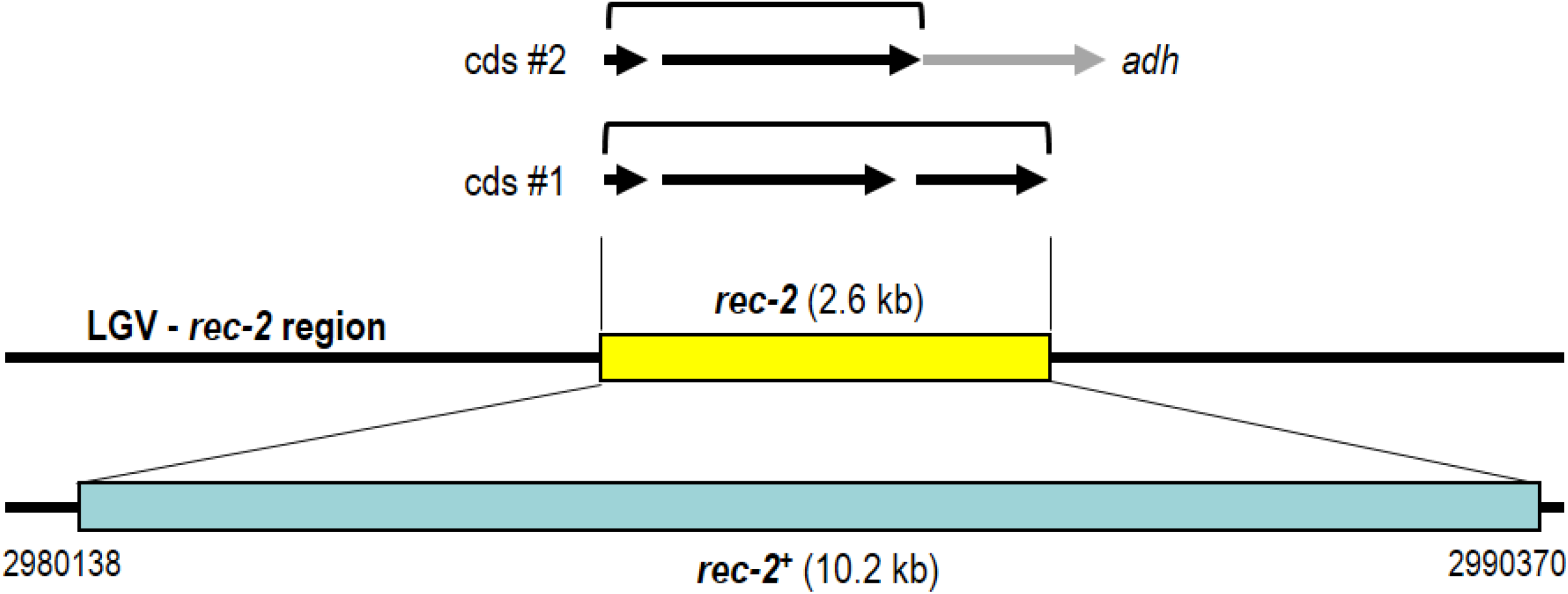
*rec-2* idiomorphs have entirely strain-specific sequences. A 10.3 kb region (light blue bar) in a *rec-2SL* strain is replaced in *rec-2* strains by a 2.6 kb fragment (yellow bar) that includes predicted coding regions: an incomplete alcohol dehydrogenase gene (*adh*), and regions (cds #1, cds #2) with homology to hypothetical proteins found in a range of filamentous fungi.

Using the predicted 762 amino acid protein to search (BLASTP 2.2.32; (ALTSCHUL *et al*. 1990; ALTSCHUL *et al*. 1997b)) non-redundant protein databases, yields substantial matches only to the carboxyl-terminal of the protein, all of which are to putative alcohol dehydrogenases (*adh* in Fig 3). In addition, a BLASTX search (ALTSCHUL *et al*. 1990; ALTSCHUL *et al*. 1997b) including sequences flanking the *rec-2LG*–specific insertion suggested that the alcohol dehydrogenase extends outside the insertion, making it an unlikely candidate for *rec-2*.

An alternate Augustus (STANKE *et al*. 2004) prediction in which the third exon is excluded yields a 552 amino acid protein (cds#2 in Fig 3; GenBank accession number BankIt2637266 rec-2 OP763471). A tblastn search (ALTSCHUL *et al*. 1997a) resulted in the highest matches (8e-20) to marine viruses, but filtering to remove query cover of less than 29% yielded similarities to only hypothetical proteins in a range of filamentous fungi. The closest match (5e-19) is to accession XM_043168630.1, an uncharacterized *Diaporthe citri* protein. There are various matches to characterized proteins, including to XM_008081332.1, a *Glarea lozoyensis* protein (CHEN *et al*. 2013) with a C2H2 and C2HC zinc finger region to which the putative REC-2 protein shows incomplete homology, XM_008085141.1, another *G. lozoyensis* gene for glutathione synthetase ATP-binding protein and AY324744.1, a *Stemphylium trifolii* elongation factor alpha gene. As the only conserved domain found in the predicted REC-2 protein is to a TolA domain, which is a membrane protein and appears unlikely to have a role in recombination, we cannot suggest a function for the transcript. Regardless of the mechanism, the REC-2 protein is clearly different from vertebrate PRDM9 (BAUDAT *et al*. 2010; MYERS *et al*. 2010; PARVANOV *et al*. 2010; BAKER *et al*. 2017).

### Mutation of *rec-2* sequence generates novel *rec-2* alleles

We attempted to replace *rec-2SL* DNA with *rec-2LG* sequence by transformation of the *hph*::*rec-2* constructs into *rec-2SL* strains of different mating types and *his-3* alleles (*his-3*^K480^ or *his-3*^K874^). In crosses between subsequent monokaryotic Hyg^R^ transformants, the frequency of His^+^ spores was expected to increase compared to the untransformed controls, which would indicate a successful replacement. Although seven transformants (three *a his-3*^K480^ and four *A his-3*^K874^) appeared to have the expected structure when subjected to Southern analysis, we saw no increase in His^+^ frequency in any cross between these transformants, suggesting that undetected ectopic duplications might exist in the transformants, resulting in RIP (SELKER 1990b).

We were able to separate transformed chromosomes from duplicated sequences in a series of crosses (Fig 4). Very low His^+^ frequencies were seen in all crosses between *his-3*^K480^ transformants and *rec-2LG his-3*^K874^ strains (Table S4). While most crosses between *his-3*^K874^ transformants and *rec-2LG his-3*^K1201^ strains were similar, with His^+^ frequencies of ~10/10^5^ viable spores (Table 3; Table S3; Fig 4), the His^+^ frequency was higher in one cross, with ~80/10^5^ viable spores (see Materials and Methods for details). Hyg^R^ progeny (T(H1); Fig 4) were extracted from this cross to select for the presence of the construct, while allowing ectopic copies to be removed by segregation, and of both mating types so they could be crossed together (Fig 4).

**Fig 4.**
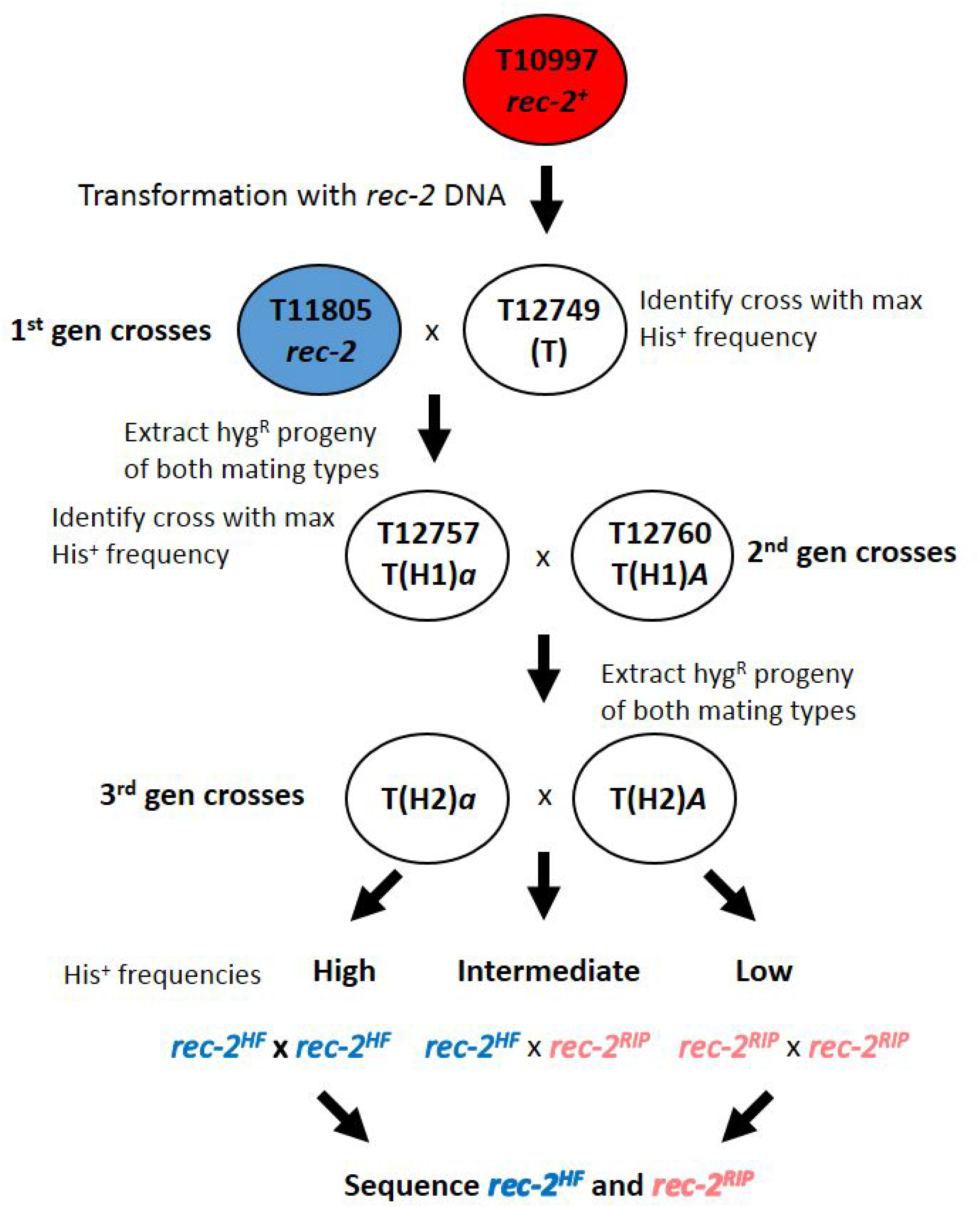
Procedure to extract novel alleles of *rec-2* from transformants. The red oval at the top of the figure represents the *rec-2^+^* strain (T10997; Table S2) transformed with the *rec-2*–replacement constructs. The first generation crosses were between homokaryons of the original transformants (T12749-T12753), represented as a white oval, and *rec-2* tester strain T11805 (Table S2), represented by a blue oval. “T” indicates the strain (T12749) that, when crossed to T11805, gave the highest frequency of His^+^ spores. For the second generation crosses, “T(H1)” represents hygromycin-resistant progeny of T12749 and T11805, of *his-3^K1201^ a* genotype (T12758-T12760) and *his-3^K874^ A* genotype (T12755-T12757). “T(H1)*a*” and “T(H1)*A*” indicate the strains (T12757 and T12760) that, when crossed together, gave the highest frequency of His^+^ spores. For the third generation crosses, “T(H2)” represents hygromycin-resistant progeny of T12757 and T12760 (T12762-T12771; Table S2).

One second generation cross (Fig 4) gave a His^+^ frequency of ~360/10^5^ viable spores (Table S3), close to half that of a cross homozygous for *rec-2LG* (YEADON *et al*. 2004), so Hyg^R^ progeny of both mating types were extracted from this cross (Fig 4). In crosses between these strains (third generation crosses; Fig 4), *rec-2* alleles were clearly segregating, giving high, intermediate or low His^+^ frequencies (Fig 4, Table 4, Table S4).

**Table 3.**
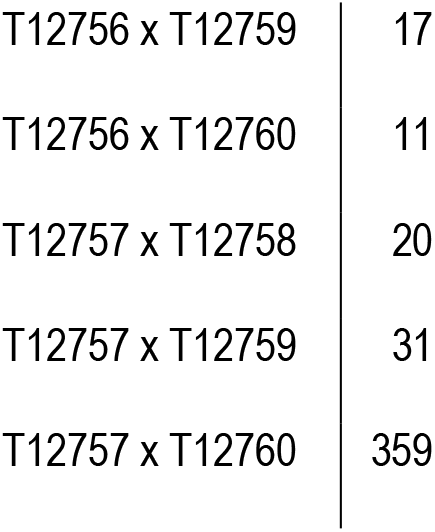
His^+^ frequency in *rec-2LG* replacement crosses. First generation crosses were between original transformants of the *his-3^K874^ rec-2SL* strain T10997 (T12749, T12750, T12751, T12752 and T12753; Table S3) and the *his-3^K1201^ rec-2LG* strain T11805 (Table S3). Progeny extracted from the cross with the highest His^+^ frequency (T12749 × T11805) were used for the 2^nd^ generation cross analyses. His^+^ frequency is per 10^5^ viable spores. Detailed cross data are provided in Table S2.

**Table 4.** Recombination phenotype segregates in the third generation progeny of *rec-2* transformants. Progeny extracted from the second generation cross with the highest His^+^ frequency were used for the third generation analysis. Both His^+^ and Lys^+^ Ad^+^ frequencies fall into three non-overlapping ranges, suggesting two alleles of *rec-2* are segregating, a high frequency *rec-2* allele (*rec-2^HF^*) and a recessive loss-of-function *rec-2* mutant (*rec-2^RIP^*). Lys^+^ Ad^+^ frequency is expressed as a percentage of the total number of viable spores, while His^+^ frequency is given per 10^5^ viable spores. Detailed cross data are provided in Table S5. Neurospora strains used for this work are listed in Table S2.

| Cross | His<sup>+</sup> ± SD | Lys<sup>+</sup> Ad<sup>+</sup> ± SD |
| --- | --- | --- |
| *rec-2<sup>HF</sup>/rec-2<sup>HF</sup>* | 1359 ± 165 | 8.4 ± 0.6 |
| *rec-2<sup>HF</sup>/rec-2<sup>RIP</sup>* | 376 ± 106 | 4.4 ± 0.5 |
| *rec-2<sup>RIP</sup>/rec-2<sup>RIP</sup>* | 15 ± 3 | 0.7 ± 0.2 |

### Sequence analysis of novel *rec-2* alleles

We sequenced the *rec-2* region in one strain (T12762 and T12763) of each allele (*rec-2^HF^* and *rec-2^RIP^* respectively), generating two sequences of approximately 3550 bases. Although this tactic (see Materials and Methods) would yield sequence from any copy in the genome, the sequence obtained is completely clean, suggesting only one copy remained. In both strains, the sequence includes 2725 bases of Neurospora *rec-2LG* region sequence followed by the *hph* sequence. Both allele sequences show multiple GC→ AT mutations, evidence of RIP (SELKER 1990a). The mutant *rec-2^RIP^* allele (GenBank accession number BankIt2637266 1c_2 OP763473) has 63 RIP mutations, generating four stop codons within cds#2 (Fig S7) and a truncated predicted protein of 212 residues. Although the sequence of the high frequency recombination *rec-2^HF^*allele (GenBank accession number BankIt2637266 1c_1 OP763472) has 26 RIP mutations, none is a stop codon so the protein is the same length as the original predicted REC-2 protein (552 residues). Of the 15 mutations within cds#2 (Fig S7), eight are synonymous, leaving seven amino acid substitutions, five of which are conservative (M30I, A402T, A406T, M407I, R410Q). The two non-conservative substitutions are near to each other (C323Y and G326R), suggesting that this section of the protein is not critical to function. Interestingly, all mutations in *rec-2^HF^* are G→A, indicating a single round of RIP, while in *rec-2^RIP^* there are 48 C→T and 15 G→A mutations, suggesting that sequences in the latter strain experienced at least two rounds of RIP (WATTERS *et al*. 1999). Alignments of the genomic sequences and the predicted coding sequences are supplied in the supplementary material (Fig S7).

## Discussion

The Neurospora *rec^+^* genes, thought to be dominant suppressors of recombination (CATCHESIDE *et al*. 1964; SMITH 1966; SMITH 1968; ANGEL *et al*. 1970; CATCHESIDE AND CORCORAN 1973; CATCHESIDE AND ANGEL 1974; CATCHESIDE 1975; CATCHESIDE 1977; CATCHESIDE 1979; BOWRING AND CATCHESIDE 1993; BOWRING AND CATCHESIDE 1999), are instead inactive idiomorphs of a recombination activator gene. Since idiomorphs lack homology, the active gene in a *rec* gene heterozygote lacks a pairing partner during meiosis and so is silenced (SHIU *et al*. 2001). The first *N. crassa* strain sequenced (GALAGAN *et al*. 2003), 74-OR23-1VA, appears to lack active sequences at all three of the known *rec* gene loci. Although regional regulation of recombination in Neurospora was reported in 1964 (CATCHESIDE *et al*. 1964), it is only now, with the advent of genome sequencing, gene replacement by transformation in *N. crassa*, and knowledge of meiotic silencing (SHIU *et al*. 2001), that the molecular basis can begin to be described.

The Neurospora predicted REC-2 protein lacks any similarity to PRDM9 and has only slight homology to other proteins in the NCBI databases, of which arguably the best match is to a *Glarea. lozoyensis* protein predicted to bind DNA. Although the 74-OR23-1VA *N. crassa* genome (GALAGAN *et al*. 2003) holds no sequences homologous to *rec-2*, a Blastp of the *Neurospora tetrasperma* and *Neurospora discreta* genomes at the Joint Genome Institute portal (http://genome.jgi.doe.gov/; (NORDBERG *et al*. 2014)) indicates good homology to several predicted proteins, with e-values of 1.7 × 10^−41^ to 1.2 × 10^−8^. The proteins match only in the central region of REC-2 (residues 120-340).

The attempt to replace *rec-2SL* DNA with the equivalent *rec-2LG* sequence presumably resulted in a transformant with more than one copy of *rec-2LG* DNA that we subsequently mutated by RIP (CAMBARERI *et al*. 1990; SELKER 1990a) to generate two novel *rec-2* alleles. We now have a mutant allele, *rec-2^HF^*, which appears to behave like wild type and another, *rec-2^RIP^*, which is completely non-functional but does not trigger meiotic silencing in a heterozygous cross due to adequate similarity with *rec-2LG*, and so has a recessive loss-of-function phenotype.

*Sad-1* increases recombination in a *rec-2LG/rec-2SL* heterozygote, but to less than half the frequency in a *rec-2LG* homozygote (Table 1), suggesting that *rec-2LG* has a dosage effect. In contrast, at both *rec-1* and *rec-3*, *Sad-1* increases recombination in *rec* heterozygotes to the same level as active allele homozygotes (Table 2), suggesting dosage may be unimportant for these mechanisms. However, it is currently unknown if the REC-1 and REC-3 proteins share homology with REC-2, which might assist in determining the mechanism by which the REC-2 protein acts and how it may differ from the other known *rec* genes.

It may have been pure chance that the three *rec* genes discovered in Neurospora all appeared to be dominant suppressors. It may also be because it is easier to detect a dominant than a recessive allele in a diploid, and all genes required for meiosis in Neurospora are essentially acting in a diploid. However, we now know that “alleles” at the *rec-1*, *rec-2* and *rec-3* loci are in reality non-homologous sequences at the same location, making them idiomorphs rather than alleles, and the dominance of one allele is due to meiotic silencing (SHIU *et al*. 2001; SHIU AND METZENBERG 2002) of the active stimulator of recombination. It is possible that yet to be discovered recombination regulation genes in Neurospora may not be subject to meiotic silencing. Nonetheless, it is interesting to note that meiotic silencing by unpaired DNA (SHIU *et al*. 2001) has been co-opted to regulate a mechanism as important to evolution as recombination hotspot activity. To our knowledge (and R.L. Metzenberg, personal communication to DC), this is the first documented biological role for meiotic silencing other than control of selfish DNA. In addition, it is clear that *rec* genes provide a method to regulate meiotic recombination additional to PRDM9 and to the default targeting to promoters seen in organisms lacking PRDM9.

## Supporting Information Legends

**Table S1: Recombination frequency in crosses between *his-3*, *am* or *his-1* alleles with and without meiotic silencing.** Strain genotypes are listed in Table S2. Spore suspensions were plated on selective media (lacking either histidine or alanine, depending on the alleles tested) and “Select” is the colony count on all five selective plates. Following dilution (800-fold for *his-3*^K874^ × *his-3*^K1201^ crosses and 400-fold for the other crosses), spore suspensions were plated on fully supplemented media. “Viable” is the colony count on all three fully supplemented plates. “Prot/10^5^” is the frequency of prototrophic progeny (those that will grow without supplementation) per 10^5^ viable spores.

**Table S2: Neurospora strains.** Loci are separated by a comma and are ordered from the left to the right arm of each linkage group. Linkage groups are separated by a semi colon and are ordered from LGI to LGVII. T12322 was obtained from Dave Jacobsen in March 2005 and is known as DJ2040-6.

**Table S3: Recombination frequency in crosses used to identify novel *rec-2* alleles, first and second generations.** Strain genotypes are listed in Table S2. Spore suspensions were poured on three selective plates containing media lacking histidine, and “Select” is the total colony count on all three plates. Following 400-fold dilution, spore suspensions were plated on fully supplemented media. “Viable” is the total colony count on the fully supplemented plates. “Prot/10^5^” is the frequency of prototrophic progeny (those that will grow without supplementation) per 10^5^ viable spores. A dash signifies an uncounted plate.

**Table S4: Recombination frequency in crosses used to identify novel *rec-2* alleles, third generation.** Strain genotypes are listed in Table S3, but all crosses are of the type *lys-4^+^ his-3^K874^ ad-3* × *lys-4 his-3^K1201^ ad-3^+^*. Spore suspensions were poured on three selective plates containing media lacking histidine, and “Select-A” is the total colony count on all three plates. Following 10-fold dilution, spore suspensions were plated on media including histidine but lacking adenosine and lysine; “Select-B” is the total colony count on these three plates. Following a further 10-fold dilution, spore suspensions were plated on fully supplemented media. “Viable” is the total colony count on the fully supplemented plates. “Prot/10^5^” is the frequency of prototrophic progeny (those that will grow without histidine) per 10^5^ viable spores. “RF%” is the frequency of progeny that will grow without adenine or lysine.

**Table S5: Primer pairs used for the *rec-2* sequence walk.** Coordinates are with reference to NC12 supercontig 5. Please note that the “forward” primers (A2F….BE17F) are reverse and the “reverse” primers (A2R….BE17R) are forward with respect to the genome sequence (GALAGAN *et al*. 2003). The 69 kb region covered by cosmids G4:D1 and X15:H7 (ORBACH 1994) begins at position 3010135 and ends at position 2940768 (Fig 2).

**Table S6: Recombination frequency in crosses including *rec-2* deletion strains.** Strain genotypes are listed in Table S2. Spore suspensions were poured on three selective plates containing media lacking histidine, and “Select” is the total colony count on all three plates. Following 800-fold dilution, spore suspensions were plated on fully supplemented media. “Viable” is the total colony count on the fully supplemented plates. “Prot/10^5^” is the frequency of prototrophic progeny (those that will grow without supplementation) per 10^5^ viable spores. Note: For control cross T10998 x T11805, spores were diluted 100-fold.

**Figure S7: CLUSTAL multiple sequence alignments.**

A: Alignment of *rec-2* allele sequences. 2.7 kb genomic sequence from *rec-2*, *rec-2^HF^* and *rec-2^RIP^* strains. Cyan highlighting indicates sequences present in *Neurospora crassa* NC12 (genomic sequence). Non-highlighted sequence (the insertion with respect to the 74-OR23-1VA sequence) begins immediately after position NC12:V: 2990402 and ends before NC12:V: 2980137, between which lie 10,265 bases in the published genomic sequence. The start codon is highlighted in green, the stop codon in red and intron sequences are in red text.

Deep pink highlighting is of RIP events where C has been replaced by T, while yellow highlighting is of G replaced by A, indicating that RIP occurred in the other strand and so in a separate event.

**B: Alignment of predicted *rec-2* amino acid sequences.** Coding sequence from *rec-2*, *rec-2^HF^* and *rec-2^RIP^* strains was translated and aligned. Yellow highlighting indicates conservative amino acid changes, cyan highlighting non-conservative changes and red highlighting novel stop codons resulting from RIP.

## Acknowledgements

This research was funded partially by the Australian Government through the Australian Research Council.

